# Immortalization and transformation of primary cells mediated by engineered ecDNAs

**DOI:** 10.1101/2023.06.25.546239

**Authors:** Davide Pradella, Minsi Zhang, Rui Gao, Melissa A. Yao, Katarzyna M. Gluchowska, Ylenia Cendon Florez, Tanmay Mishra, Gaspare La Rocca, Moritz Weigl, Ziqi Jiao, Hieu H.M. Nguyen, Felix Grimm, Marta Lisi, Chiara Mastroleo, Kevin Chen, Jens Luebeck, Vineet Bafna, Cristina R. Antonescu, Andrea Ventura

## Abstract

Focal gene amplifications are among the most common cancer-associated mutations, but their evolution and contribution to tumorigenesis have proven challenging to recapitulate in primary cells and model organisms. Here we describe a general approach to engineer large (>1 Mbp) focal amplifications mediated by extrachromosomal circular DNAs (ecDNAs, also known as “double minutes”) in a spatiotemporally controlled manner in cancer cell lines and in primary cells derived from genetically engineered mice. With this strategy, ecDNA formation can be coupled with expression of fluorescent reporters or other selectable markers to enable the identification and tracking of ecDNA-containing cells. We demonstrate the feasibility of this approach by engineering MDM2-containing ecDNAs in near-diploid human cells, showing that GFP expression can be used to track ecDNA dynamics under physiological conditions or in the presence of specific selective pressures. We also apply this approach to generate mice harboring inducible *Myc*- and *Mdm2*-containing ecDNAs analogous to those spontaneously occurring in human cancers. We show that the engineered ecDNAs rapidly accumulate in primary cells derived from these animals, promoting proliferation, immortalization, and transformation.

## MAIN

Increased oncogene expression mediated by gene amplifications is common in human cancers ^1^. Two major classes of amplification have been described: chromosomal and non-chromosomal. The latter is characterized by the presence of multiple copies of circular DNAs that are thought to originate from the fragmentation and subsequent circularization of segments of chromosomes ^2, 3^. These large (0.5-3 Mbp) “extrachromosomal circular DNAs” (ecDNAs) are also known as “double minutes” for their paired appearance in metaphase spreads ^4^ (reviewed in ^5–8^).

Analyses of The Cancer Genome Atlas (TCGA) and the International Cancer Genome Consortium (ICGC) databases show that ecDNAs are highly prevalent across multiple cancer types and are associated with unfavorable prognosis ^9, 10^. At least two dozen oncogenes have been reported in circular amplicons, with *MDM2, EGFR,* and *MYC* being the three most common ^9^.

EcDNAs are advantageous to cancer cells for several reasons. First, ecDNAs lack centromeric sequences and therefore segregate randomly during mitosis ^11,12, 13^. This enables the accumulation of large numbers of ecDNAs, thus promoting intra-tumoral heterogeneity and accelerating tumor evolution^9^. The heterogeneity in copy number and the speed with which ecDNAs can be gained and lost allow tumor cells to adapt to changes in the tumor microenvironment. For example, the reversible loss of EGFR-containing ecDNAs in glioblastoma cells in response to EGFR inhibitors is an important mechanism underlying drug resistance ^14^. Genes residing on ecDNAs are also transcribed more efficiently than when residing on a linear chromosome, likely due to a more accessible chromatin configuration and the lack of higher-order compaction ^7, 15^. Finally, ecDNAs can directly affect the expression of other cancer-promoting genes by acting as mobile enhancers, regulating gene expression *in trans* ^12, 16^.

Many fundamental questions about ecDNA biology remain unanswered. For example, despite recent reports showing the presence of ecDNAs in precancerous esophageal lesions ^17^, the exact roles of oncogene-containing ecDNAs in tumor initiation and progression are poorly understood. It is also unclear whether primary cells possess mechanisms to protect against the formation or propagation of ecDNAs, and whether the presence of ecDNAs exposes novel therapeutically actionable vulnerabilities. The ability to induce and track the formation of specific ecDNAs in primary cells and whole organisms, with precise spatiotemporal control, would greatly accelerate progress in this field. Here we describe a novel general strategy that addresses these issues, and we apply it to engineer the formation of oncogene-containing ecDNAs in cells and mice.

### An inducible system to model ecDNA formation

To develop a flexible system to induce the formation of ecDNA analogous in size and behavior to those found in human cancers, we took advantage of the ability of the Cre recombinase to induce circularization of any region flanked by two *loxP* sites having the same orientation ^18, 19^. We reasoned that by flanking a region of interest in the human or murine genome with *loxP* sites, it should be possible to induce the generation of circular DNA molecules that recapitulate ecDNAs detected in human cancer cells (Fig. 1A).

**Figure 1.**
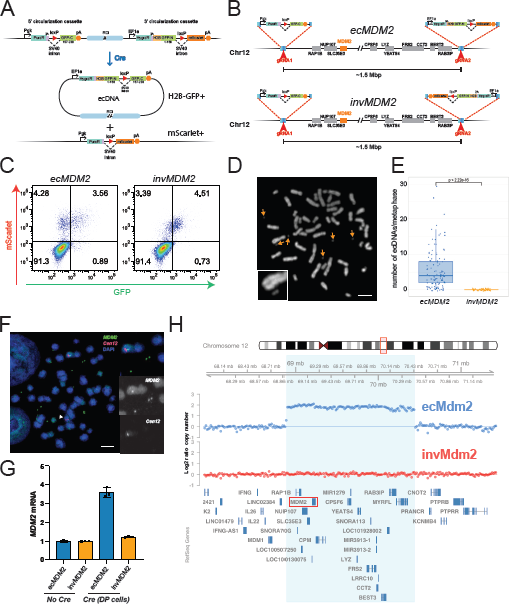
A general strategy for ecDNA engineering. **A.** Schematic of the circularization strategy. ROI = region of interest. **B.** Schematics of the *ecMDM2* and *invMDM2* alleles generated in HCT116 cells. The circularization cassettes are not drawn to scale. Notice that in the *invMDM2* allele the 3’ cassette is inserted in the opposite orientation to induce inversion rather than circularization. **C.** Flow cytometry scatter plot of representative *ecMDM2* and *invMDM2* clones 6 days after AdCre infection. **D.** Representative metaphase spread obtained from sorted double positive ecMDM2 cells. Orange arrows indicate double minutes/ecDNAs. Inset, shows a magnified ecDNA next to a chromosome. scale bar = 5µm **E.** Box-and-whiskers showing the distribution of the number of ecDNAs observed in metaphases from double positive *ecMDM2* and *invMDM2* cells (p-value = two-tailed Fisher t-test, more than 50 metaphases per genotype were analyzed). **F.** Representative DNA-FISH using *Mdm2* (green) and chromosome 12 centromere (red) probes performed on metaphase spreads from double positive *ecMDM2* cells. Notice the presence of numerous *MDM2*-positive double minutes (insets), and the concomitant loss of *MDM2* signal from one of the two chromosome 12 (white arrowhead). Scale bar: 5µm. **G.** Relative *MDM2* mRNA abundance as determined by RT-qPCR in control and sorted double positive cells AdCre infected *ecMDM2* and *invMDM2* cells. **H.** Genomic DNA from sorted double positive *ecMDM2* and *invMDM2* cells was analyzed by shallow whole genome sequencing and the log2 ratio of relative copy number across the entire 15kb bins was determined using the QDNAseq R package. The panel shows relative copy number across the region of chromosome 12 surrounding the region flanked by the two circularization cassettes (light blue). Notice the presence of a focal amplification exactly matching the predicted ecDNA boundaries, only in *ecMDM2* cells.

To facilitate the identification and selection of cells harboring the engineered ecDNAs, we designed two “circularization cassettes” to be inserted *in cis* at the desired circularization breakpoints (Fig. 1A). Cre-mediated recombination between the two *loxP* sites contained in the cassettes will lead to circularization of the intervening genomic region and reconstitution of a functional H2B-GFP transgene that is encoded by the resulting ecDNA (Fig. 1A **and** Supp. Fig. 1A). The 3’ cassette contains a second fluorescence marker (mScarlet) whose expression is also induced upon Cre-expression but remains on the linear chromosome. This dual color system allows discrimination between cells that have undergone circularization but have subsequently lost the ecDNA(s) due to random segregation (GFP-;mScarlet+) (Supp. Fig. 1A), and cells harboring a tandem duplication caused by Cre-mediated recombination between sister chromatids (GFP+;mScarlet-). Finally, the hygromycin resistance gene is included in the ecDNA upon Cre-mediated circularization and can be used to modulate ecDNA abundance.

To test this strategy, we chose the HCT116 colorectal cancer cell line, which is near-diploid, chromosomally stable, and harbors no endogenous ecDNAs. We used CRISPR-Cas9-mediated genome editing to insert the two circularization cassettes flanking an approximately 1.5 Mbp region containing, among other genes, the human *MDM2* locus (Fig. 1B). The *MDM2* oncogene, which was originally cloned from a cell line containing double minutes^20^, encodes for a key negative regulator of the p53 pathway^21, 22^ and is frequently amplified in ecDNAs in sarcomas and other human cancers ^23–25^.

As a control we generated a companion HCT116 line in which the 3’ cassette is inserted at the same location, but in the opposite orientation (Fig. 1B). With this configuration, the genomic region flanked by the two cassettes is inverted rather than excised upon Cre-expression, and therefore both fluorescent reporters remain linked to the linear chromosome (Supp. Fig. 1B). We refer to these two engineered cell lines as “*ecMDM2*” and “*invMDM2*”, respectively.

After selection with puromycin and hygromycin, individual clones were genotyped and sequenced to identify those with correct integration of both cassettes in the correct orientation. Successfully targeted *ecMDM2* and *invMDM2* clones were expanded and infected with Cre-expressing recombinant adenoviruses (AdCre) to induce circularization or inversion, respectively. Five to six days after infection, each clone was examined by flow cytometry. Two classes of clones were identified: one with low (<1%) and one with higher (>5%) frequency of mScarlet+ cells (Fig. 1C and Supp. Fig. 1C). Since the efficiency of Cre-mediated recombination depends on the distance between the two *loxP* sites, we concluded that the two circularization cassettes were targeted *in cis* in the clones with higher frequency of mScarlet+ cells.

To test whether GFP expression in *ecMDM2* cells reflects the formation of ecDNAs, we generated metaphase spreads from sorted double positive (GFP^+^;mScarlet^+^) *ecMDM2* and *invMDM2* cells. Most metaphases from e*cMDM2* double positive cells—but none from *invMDM2* cells—contained *MDM2*-positive double minutes. As expected, we observed concomitant loss of *MDM2* signal from one of the two copies of chromosome 12 in metaphases from *ecMDM2* cells (98% of metaphases examined), but not in metaphases from *invMDM2* cells (Fig. 1D-F). Consistent with this observation, sorted double positive *ecMDM2* cells had higher *MDM2* mRNA levels compared to control (no Cre) and sorted double positive *invMDM2* cells (Fig. 1G). Furthermore, shallow whole genome sequencing (sWGS) of sorted double positive cells showed the presence of a focal amplification mapping precisely to the 1.5 Mbp region flanked by the two circularization cassettes in *ecMDM2*, but not in *invMDM2* cells (Fig. 1H). These results demonstrate that the strategy we have developed can be used to model the formation and propagation of large ecDNAs in cells.

### Tracking ecDNA abundance and dynamics in live cells

To determine whether GFP expression could be used to track ecDNA abundance and dynamics, we generated metaphase spreads from sorted GFP+;mScarlet+ *ecMDM2* cells into three bins based on GFP intensity and analyzed them (Fig. 2A). The average number of ecDNAs per cell correlated closely with GFP intensity (Fig. 2B) and this positive correlation between GFP expression and *MDM2* ecDNA abundance was further confirmed by sWGS (Fig. 2C).

**Figure 2.**
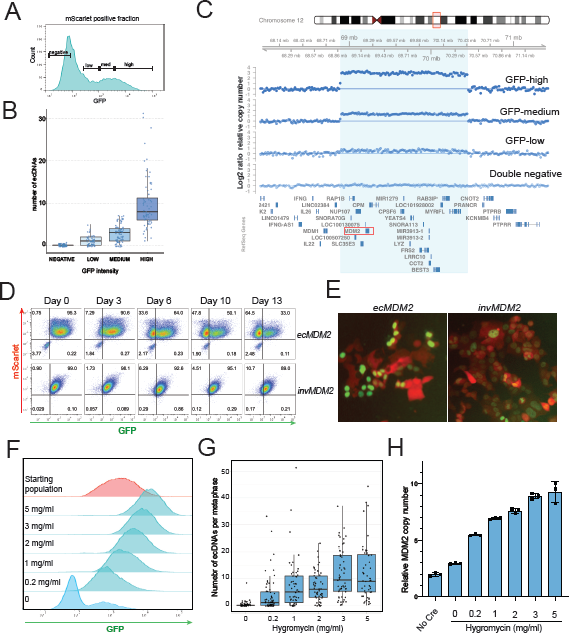
Dynamics of engineered ecDNAs in HCT116 cells. **A.** Sorted double positive cells from an AdCre-infected *ecMDM2* line were expanded in culture in the absence of hygromycin. The mScarlet+ population was sorted into 4 bins based on GFP intensity (negative, low, medium, and high), as indicated in the histogram plot. **B.** Box-and-whiskers plot showing the number of ecDNAs per metaphase in the sorted population described in panel A. **C.** *ecMDM2* cells were infected with AdCre, sorted according to mScarlet and GFP intensity and analyzed by shallow whole genome sequencing. The panel shows log2 ratio of relative copy number across the region of chromosome 12 surrounding the predicted ecDNA (defined by the light blue highlight). **D.** Sorted double positive cells from AdCre infected *ecMDM2* and *invMDM2* clones were cultured in the absence of hygromycin and analyzed by flow cytometry at the indicated time points. **E.** Representative fluorescence microscopy images from double positive sorted *ecMDM2* and *invMDM2* cells maintained in media without hygromycin for ∼ 1 week. **F.** AdCre infected and sorted double positive *ecMDM2* cells were expanded for 13 days in medium containing the indicated concentration of hygromycin and analyzed by flow cytometry. Histogram plot of GFP fluorescence is shown. **G.** Box-and-whiskers plot showing the number of ecDNAs per metaphase observed in the cells described in panel E. **H.** *MDM2* copy number as determined by qPCR in cells from panel F.

Having established that GFP intensity can be used as a surrogate for ecDNA presence and abundance, we next examined the dynamics of the engineered *MDM2*-containing ecDNAs in HCT116 cells. We infected *ecMDM2* and *invMDM2* cells with AdCre and measured the fraction of GFP+, mScarlet+, and GFP+;mScarlet+ cells at different time points. For both *ecMDM2* and *invMDM2* cells, the fraction of double positive cells peaked at approximately 5-6 days post infection (Supp. Fig. 2A). However, while the double positive population in *invMDM2* cells remained roughly constant at later time points, in *ecMDM2* cells we observed a marked reduction of this cell population with concomitant accumulation of GFP-;mScarlet+ cells.

To test whether the GFP-;mScarlet+ cells derive from initially double positive cells, we sorted double positive cells from *ecMDM2* and *invMDM2* cells and serially passaged them. Again, *invMDM2* cells remained largely double positive throughout the experiment, but *ecMDM2* cells gradually lost GFP expression (Fig. 2D-E). These results demonstrate that analogously to ecDNAs observed in human cancers, the engineered ecDNAs segregate independently from the linear chromosome from which they have originated. Furthermore, the observation that *ecMDM2* cells gradually lose ecDNAs indicate that extra copies of the engineered ecDNAs may not provide a fitness advantage to HCT116 cancer cells *in vitro*^26^.

To determine whether an artificially imposed selective pressure would promote the accumulation of *MDM2-*containing ecDNAs, we took advantage of the presence of the hygromycin resistance gene in the engineered ecDNAs (Fig. 1A **and** Supp. Figure 1A). When sorted double positive *ecMDM2* cells were cultured in the presence of increasing concentrations of hygromycin we observed a proportional increase in GFP intensity in the population (Fig. 2F **and** Supp. Fig. 2B). This was accompanied by an increase in *MDM2* copy number (Fig. 2G) and a concomitant accumulation of *MDM2*-containing ecDNAs (Fig. 2H **and** Supp. Figure 2C). Analogous results were obtained when freshly AdCre-infected *ecMDM2* cells were maintained in the presence of high hygromycin concentration (not shown).

While analyzing these experiments we also noticed the presence of homogeneously staining regions (HSR) decorated by the *MDM2* probe instead of double minutes in approximately 6% of metaphases generated from double positive sorted *ecMDM2* cells (Supp. Fig. 2E). In all cases, one of the two chromosome 12 showed loss of endogenous *MDM2* staining, confirming that these cells had successfully circularized and excised the *ecMDM2* allele upon Cre expression. This indicates that the engineered ecDNA can occasionally re-integrate into a chromosome, recapitulating a poorly understood phenomenon that has been previously reported for naturally occurring ecDNA ^2, 3, 27^.

Together, these results demonstrate the successful engineering and non-invasive tracking of functional and replication-competent oncogene-containing ecDNAs in human cells.

### Generation of genetically engineered mice with inducible ecDNAs

We next sought to determine whether a similar strategy could be adapted to engineer ecDNA formation in primary cells and in mice. We employed CRISPR-Cas9 based genome editing by zygote electroporation^27^ to generate mice harboring *loxP* sites spanning a 1.7 Mbp genomic region on chromosome 15 containing the *Myc* gene, an oncogene that is amongst the most commonly amplified in ecDNAs in human cancers ^9, 10^ (Fig. 3A **and** Supp. Fig. 3A).

**Figure 3.**
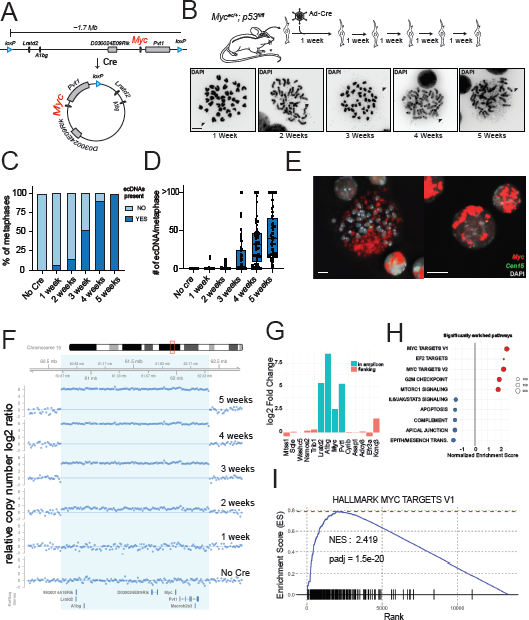
Generation of Myc-containing ecDNAs in primary cells from genetically engineered mice. **A**. Schematic of the *Myc^ec^* allele and of the ecDNA generated upon Cre-mediated recombination. **B**. AdCre-treated aNSC derived from *Myc^ec/+^; p53^fl/fl^* mice were propagated *in vitro*. Metaphase spreads (lower panels) at different time-points were collected and representative micrographs are shown. Arrowheads indicate double minutes/ecDNAs. Scale bar: 5 µm. **C**. Barplot showing the fraction of metaphases with or without ecDNAs and (**D**), box-and-whiskers plot showing the number of ecDNA per metaphase in *Myc^ec/+^; p53^fl/fl^* aNSC at different times upon AdCre infection. Cells not infected with AdCre are also shown. **E**. metaphase (left) and interphase FISH (right) performed on aNSC from *Myc^ec/+^; p53^fl/fl^* 5 weeks upon AdCre infection. The *Myc* signal is shown in red. Control probe labeling a pericentromeric region of chromosome 15 is in green. Scale bar: 5 µm. **F**. Shallow whole genome sequencing analysis of *Myc^ec/+^; p53fl/fl* aNSC collected at different time-points post AdCre infection. The panel shows relative copy number across the region of chromosome 15 surrounding the predicted *Myc*-containing ecDNA (light blue). Notice the progressive increase in copy number of a focal amplification exactly matching the predicted ecDNA boundaries. **G**. Log2 fold change of mRNA expression of genes included (blue) or flanking (red) the engineered *Myc* amplicon in *Myc^ec/+^; p53^fl/fl^* vs *p53^fl/fl^* aNSC 5 weeks upon AdCre infection, as determined by RNAseq analysis. **H**. Gene set enrichment analysis showing the top enriched Hallmark pathways in *Myc^ec/+^;p53^fl/fl^* vs *p53^fl/fl^* aNSC at 5 weeks after AdCre infection. **I**. Gene set enrichment plot of Hallmark gene set MYC Targets V1, the most enriched gene set in *Myc^ec/+^; p53^fl/fl^* aNSC. NES: Normalized Enrichment Score.

For this set of experiments, to avoid the risk of interfering with expression of endogenous genes in the ecDNAs and to more closely recapitulate ecDNAs observed in human cancers, we only inserted the *loxP* sites rather than the full circularization cassettes described for the *in vitro* experiments.

We confirmed the insertion of *loxP* sites on chromosome 15 in the correct orientation *in cis* by Sanger sequencing and by analyzing the progeny of founder mice crossed to wild type mice, respectively (Supp. Fig. 3). This new engineered allele will be referred to hereafter as *Myc^ec^*. *Myc^ec/+^* mice were obtained at the expected Mendelian frequency, were viable and fertile, and showed no obvious developmental defect or phenotype (Supp. Fig. 3B-E).

We first attempted to induce ecDNA formation in primary cells derived from *Myc^ec^* animals. To prevent replicative and oncogene-induced senescence or apoptosis caused by increased MYC expression, we intercrossed *Myc^ec^* and *p53^fl^* mice^28^ to obtain *Myc^ec/+^; p53^fl/fl^*animals.

From these animals and from *p53^fl/fl^* littermate controls, we isolated adult neural stem cells (aNSC) and infected them with AdCre. Following infection, these cells were propagated *in vitro* and evaluated for the presence of double minutes at weekly intervals for a total of 5 weeks (Fig. 3B). At one-week post-infection, only a small fraction of *Myc^ec/+^; p53^fl/fl^* aNSCs contained relatively few ecDNAs, consistent with Cre-mediated circularization occurring only in a relatively small number of cells. Over the ensuing 4 weeks, however, the number of metaphases with ecDNAs and the number of ecDNAs per metaphase increased dramatically such that at five weeks every metaphase we examined showed multiple ecDNAs (Fig. 3C). In contrast, we did not detect ecDNAs at any time point in the control cells (data not shown). The increased abundance of ecDNAs per metaphase was accompanied by increased *Myc* copy number as determined by qPCR (Supp. Fig. 4A). Importantly, FISH analysis confirmed that all ecDNAs contained the *Myc* locus (Fig. 3E) and sWGS showed the presence of a focal amplification that mapped exactly to the region flanked by *loxP* sites (Fig. 3F). Interestingly, DNA-FISH performed on interphase nuclei showed the *Myc* signal to be often localized into discrete clusters, rather than being randomly distributed (Fig. 3E **and** Supp. Fig. 4B), in agreement with a series of recent studies performed on naturally occurring *MYC*-containing ecDNAs ^29, 30^.

**Figure 4.**
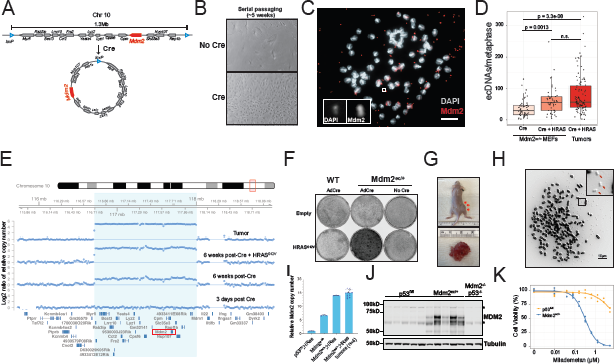
Engineered Mdm2-containing ecDNAs promote immortalization and transformation of primary mouse cells. **A.** Schematic of the *Mdm2^ec^* allele and of the ecDNA generated upon Cre-mediated recombination. **B.** Representative image of *Mdm2^ec/+^* MEFs maintained in culture for six weeks after being infected with AdCre or left untreated. **C.** Representative metaphase spread and DNA-FISH showing the presence of multiple *Mdm2*-positive ecDNAs in AdCre-infected *Mdm2^ec/+^* MEFs (inset). Scale bar = 10μm. **D.** Box-and-whiskers plot showing the number of ecDNAs per metaphase in AdCre infected *Mdm2^ec/+^*MEFs transduced or not with HRAS^G12V^ and in tumors (n = 4) derived from injecting the HRAS-transduced cells in the flank of nude mice. p-values = pairwise Wilcoxon Rank Sum Test. **E.** Shallow whole genome sequencing of AdCre-infected *Mdm2^ec/+^* MEFs at the indicated time points after AdCre infection showing progressive amplification of the region flanked by the two *loxP* sites over time. Concomitant expression of an oncogenic HRAS and results in a further increase in copy number. “Tumor” indicates sWGS of a tumor obtained by injecting the HRAS-infected cells into the flank of a nude mouse. The absence of signal in the chr10:118199900-118279700 region downstream of the amplicon is an artifact due to QDNASeq automatically removing low-mappability regions from the analysis. **F.** Focus formation assay of MEFs with the indicated genotype and Cre-treatment, transduced with either pBABE or pBABe-HRAS^G12V^. **G.** Representative image of a tumor developing in a nude mouse injected subcutaneously with 400,000 AdCre- and HRAS^G12V^-infected *Mdm2^ec/+^* MEFs. **H.** Metaphase spread showing the presence of numerous ecDNAs (arrows) in cells isolated from the tumor shown in panel G. **I.** *Mdm2* copy number by qPCR in control cells and in cells derived from the tumor in panel G. **J.** Anti-MDM2 immunoblot of lysates from tumors generated by injecting HRAS^G12V^-transduced MEFs with the indicated genotypes into nude mice. Mdm2-p53 DKO tumors were included as negative controls for the MDM2 antibody. Asterisks indicated non-specific bands**. K.** Dose-response curve of *p53^fl/fl^;HRAS^G12V^*and *Mdm2^ec/+^;HRAS^G12V^*MEFs treated with the MDM2 antagonist milademetan for 72 hours. Mdm2-p53 double knockout tumors were included as negative controls for the MDM2 antibody. Asterisks indicated non-specific bands**. K.** Dose-response curve of *p53^fl/fl^;HRAS^G12V^* and *Mdm2^ec/+^;HRAS^G12V^*MEFs treated with the MDM2 antagonist milademetan.

The rapid increase in the frequency of ecDNAs-containing cells and the accumulation of multiple copies of the *Myc*-containing ecDNAs, shows that their presence in these primary cells is well tolerated and provides a strong fitness benefit. Indeed, the accumulation of engineered ecDNAs resulted in increased *Myc* mRNAs expression and protein levels (Supp. Fig. 4D and 4E), as well as increased proliferation *in vitro* (Supp. Fig. 4F).

To gain insights into the transcriptional changes caused by the accumulation of engineered ecDNAs containing the *Myc* locus, we performed RNAseq analysis comparing *p53^fl/fl^*and *Myc^ec/+^; p53^fl/fl^* aNSCs 5 weeks after AdCre infection (Fig. 3G, Supp. Fig. 4, **and Supp. Table 1**). While *Myc^ec/+^; p53^fl/fl^* cells showed upregulation of Myc and the other genes included in the amplicon, we noticed that the *Myc* mRNA was upregulated less than expected based on the copy number change (Fig. 3G, Supp. Fig. 4G, and **Supp. Data 1**). This is consistent with previous studies showing the existence of a negative autoregulatory loop controlling expression of the endogenous *Myc* locus ^31^ and emphasizes the fundamental difference between modeling focal amplification and ectopically expressing an oncogene under the control of an artificial promoter. Finally, we confirmed global activation of the MYC-driven transcriptional program in the *Myc^ec/+^; p53^fl/fl^* aNSC by performing gene set enrichment (FIg. 3H-I and Supp. Fig. 4H).

Next, we examined whether ecDNA formation could be induced directly *in vivo*. We crossed *Myc^ec/+^* mice to *Rosa26-Cre-ER^T^*^2^ ^32^ or *Nestin-Cre* mice ^33^, to induce ecDNA formation in a temporally and spatially controlled fashion. ecDNA formation was readily detected in the brain of *Nestin-Cre* mice, and in a wide range of tissues following tamoxifen administration in *Rosa26-Cre-ER^T^*^2^ animals (Supp. Fig. 5A-B). In addition, DNA-FISH performed on the brain of *Myc^ec/+^; Rosa26-Cre-ER^T^*^2^ mice treated with tamoxifen showed the presence of a subset of cells harboring multiple copies of the *Myc* locus, consistent with accumulation of ecDNAs. Collectively, these results demonstrate the generation of functional, self-replicating, oncogene-containing ecDNAs both *ex vivo* and *in vivo* (Supp. Fig. 5C).

### Mdm2-containing engineered ecDNAs overcome p53-dependent senescence

To further explore the oncogenic potential of engineered ecDNAs, we generated a second mouse strain harboring *loxP* sites flanking a 1.3 Mb region containing the *Mdm2* oncogene and a dozen other genes (*Mdm2^ec^*, Figure 4A **and** Supp. Fig. 6).

*Mdm2^ec/+^* and *Mdm2^ec/ec^* mice are viable and fertile and do not display any obvious abnormality. To test whether induction of ecDNAs containing the *Mdm2* locus would result in *Mdm2* amplification and suppression of the p53 pathway, we serially passaged MEFs isolated from *Mdm2^ec/+^*and wild type embryos. After one passage, MEFs were either left untreated or infected with AdCre to induce circularization of the *Mdm2*-containing chromosomal region (Supp. Fig. 6E). As expected, control untreated *Mdm2^ec/+^* and wild type MEFs stopped proliferating after approximately 3 weeks in culture and acquired the characteristic morphology of senescent cells (Fig. 4B). In contrast, AdCre infected *Mdm2^ec/+^* MEFs became immortalized (Fig. 4B). Metaphase spreads generated from these cells after 6 weeks in culture revealed the presence of numerous double minutes containing the *Mdm2* locus in the vast majority of metaphase analyzed (Fig. 4C**, D**). This correlated with markedly increased *Mdm2* RNA and protein expression in these cells (Supp. Fig. 7), and shallow whole genome sequencing revealed the presence of a focal gene amplification matching exactly the region flanked by the two *loxP* sites (Fig. 4E). These results demonstrate that engineered *Mdm2*-containing ecDNAs rapidly accumulate in primary MEFs and promote their immortalization.

### Engineered Mdm2-containing ecDNAs enable oncogenic transformation of primary cells

Functional or genetic inactivation of the p53 pathway is required for transformation of MEFs by ectopically expressed oncogenic RAS^34^. To test whether engineered *Mdm2*-containing ecDNA would recapitulate the oncogenic function of naturally occurring *MDM2*-containing double minutes, we transduced early passage control and Cre-treated *Mdm2^ec/+^*MEFs with retroviruses encoding the *HRAS^G12V^* oncogene. After an initial burst of proliferation, control cells underwent cell cycle arrest and became senescent. In contrast, AdCre infected *Mdm2^ec/+^* cells continued proliferating and acquired a transformed phenotype characterized by loss of contact inhibition (Fig. 4F).

When injected subcutaneously into immunocompromised mice, *HRAS^G12V^*transduced AdCre-infected *Mdm2^ec/+^* MEFs invariably formed tumors (7/7, two independent MEF lines). In contrast, no tumors developed in mice injected with *HRAS^G12V^*-transduced wild type cells (0/5) or *HRAS^G12V^*-transduced *Mdm2^ec/+^* MEFs that had not been previously infected with AdCre (0/4). Tumors formed by the AdCre-infected *Mdm2^ec/+^*MEFs retained marked *Mdm2* amplification, invariably contained a large number of ecDNAs, and expressed high levels of *Mdm2* RNA and protein (Fig. 4G-J **and** Supp. Fig. 8A). Histological characterization of the resulting tumors showed a high-grade spindle cell sarcoma arranged in short fascicles characterized by ovoid cells with marked nuclear pleomorphism, frequent mitosis, and infiltration of the surrounding skeletal muscle (Supp. Fig. 8B-E). In addition, focal areas of the lesion displayed cells with distinct single or multi-intracytoplasmic fat vacuoles and focal nuclear indentation, a characteristic feature of lipoblasts (Supp. Fig. 8D). The alternating high grade undifferentiated component with areas with adipocytic differentiation are suggestive of a diagnosis of dedifferentiated liposarcoma, a tumor whose hallmark genetic aberration is the amplification of *MDM2*.

Finally, to confirm the causative role of *Mdm2* amplification in the immortalization and transformation of these cells, we exposed them to increasing concentrations of milademetan, an MDM2 antagonist currently in clinical trials ^34^. Milademetan potently inhibited the growth of the *HRAS^G12V^* infected *Mdm2^ec/+^* cells at nanomolar concentrations but was largely ineffective on HRAS-transformed p53-null MEFs (Fig. 4K). importantly, milademetan treatment stabilized the p53 tumor suppressor and markedly upregulated *Mdm2* and *p21-Waf1* mRNAs and protein products, two well characterized direct transcriptional targets of p53 ^35^ (Supp. Fig. 9).

Collectively, these results demonstrate that engineered *Mdm2*-containing ecDNAs rapidly accumulate in primary cells, cooperate with the Ras oncogene to promote their oncogenic transformation, and are maintained *in vivo*. They also provide direct experimental evidence that focal gene amplification mediated by ecDNAs can drive oncogenic transformation.

## Discussion

The ability to engineer cancer-associated genetic lesions in cells and in model organisms is essential to dissect their contribution to tumor initiation and progression and to develop more accurate preclinical models of human cancers. Advances in germline and somatic gene editing methods over the past few decades have enormously improved our ability to control the formation of a wide range of loss-of-function and gain-of-function mutations in a temporally and spatially controlled fashion. Despite this progress, focal amplification events, a common mechanism for oncogene activation in human cancers, had not yet been modeled. Surrogate approaches relying on ectopic expression of individual oncogenes via transgenes, while useful, fail to recapitulate the complexity, intra-tumoral heterogeneity in copy number, and evolution of naturally occurring gene amplification events.

The general strategy to engineer the formation of ecDNA amplifications described here overcomes these limitations. In the first implementation of this strategy, we generated human cancer cell lines in which *MDM2*-containing ecDNAs are formed upon Cre-expression. We show that ecDNA-containing cells can be visualized and tracked using fluorescent reporters encoded by the ecDNAs and the linear chromosome from which they originate. We apply a similar strategy to engineer *Mdm2-* and *Myc-*containing ecDNAs in mice and in cells derived from these animals, and we show that these ecDNAs confer a selective advantage to primary cells harboring them, accumulate over time *in vitro,* and promote cell proliferation and immortalization. Finally, we show that engineered *Mdm2* containing ecDNAs cooperate with oncogenic Ras to induce the oncogenic transformation of mouse fibroblasts.

We believe these results have significant implications. They demonstrate that oncogene-containing ecDNAs can be generated and rapidly accumulate in primary non-transformed cells such as MEFs and aNSCs. This is relevant because until now oncogene-containing ecDNAs had been observed only in established cancer cell lines and primary tumors, but not in normal cells. Analogously, previous attempts to induce the formation of ecDNAs using CRISPR-Cas9 ^26, 36^ or through drug selection ^3, 37^ had been performed in cancer cell lines, but not in primary cells or in whole organisms. Such approaches also do not provide a way to track ecDNA-containing cells non-invasively and have not modeled oncogenic ecDNAs.

We also expect that the general strategy we have developed and the reagents we have generated will prove useful in dissecting multiple aspects of ecDNA biology that are currently unresolved. For example, the presence of selectable markers and fluorescence reporters in the engineered ecDNAs is ideally suited to perform large scale CRISPR-screens aimed at identifying factors that are required for ecDNA propagation and maintenance, and to identify unique genetic vulnerabilities conferred to cancer cells by the presence of ecDNAs. Similarly, the presence of reporters encoded by the engineered ecDNAs provides an opportunity to test the tantalizing hypothesis that ecDNAs can be transferred horizontally, either through direct cell contact or through secreted vesicles.

The ability to induce the formation of ecDNAs in primary cells will also facilitate studies aimed at investigating the mechanisms underlying chromatin changes that have been recently described in cancer associated ecDNAs, and the role of DNA regulatory elements present in human ecDNAs that have been proposed to affect gene expression *in trans* ^29, 30^.

It is also worth noting that although for the present studies we used the Cre-lox system to induce circularization, the same results can be achieved using other site-specific recombinases (i.e. Flpe or Dre)^38^, further diversifying the potential applications of this strategy to model tumor evolution *in vivo*.

Despite these benefits, the approach described here has some limitations. First, while our strategy recapitulates the subset of ecDNAs generated by two double stranded DNA breaks followed by recircularization, ecDNAs can also result from chromothripsis, for which Cre-induced recombination is not an accurate proxy.

Second, the efficiency of Cre-induced circularization is known to decrease as the distance between the *loxP* site increases. For the ecDNAs generated in this study, we have observed an efficiency of circularization of 7-20%. The size of the ecDNAs we have engineered spans ∼1.3 Mbp to ∼1.7 Mbp, well within the range of many naturally occurring cancer-associated ecDNAs, but it is possible that circularization efficiency will be substantially lower when attempting to model larger ecDNAs (3 Mbp or more).

Third, for the *in vitro* experiments we used H2B-GFP as a reporter for the generation of ecDNAs. While H2B-GFP has the advantage of labeling the chromatin, allowing the direct visualization of ecDNAs during mitosis, GFP variants with shorter half-lives may be better suited to study ecDNA dynamic in cell populations and to facilitate pooled CRISPR-screens.

A puzzling aspect of this study is that although the engineered ecDNAs are replication competent, confer a strong selective advantage *ex vivo,* and can immortalize and transform primary cells, we have not yet observed the development of autochthonous tumors harboring amplified ecDNAs in *Myc^ec/+^* or *Mdm2^ec/+^* animals. This is not due to issues with Cre-mediated circularization *in vivo*, as we can readily detect circularization in multiple tissues upon Cre-delivery.

Several explanations are possible. First, oncogenic ecDNAs alone may not be capable of initiating tumorigenesis *in vivo*, and additional oncogenic events are required. It is also conceivable that cell autonomous or non-cell autonomous tumor suppressive mechanisms *in vivo* prevent the expansion of ecDNA-containing cells. Second, although recent work has shown the presence of ecDNAs in early pre-cancerous lesions in the esophagus^17^, it is possible that in other contexts ecDNAs are acquired relatively late in tumorigenesis and do not contribute significantly to tumor initiation. Lastly, tumor formation driven by ecDNAs could have much longer latency than tumorigenesis driven by other oncogenic events. Indeed, the nature of focal amplifications driven by ecDNAs requires multiple rounds of cell division for ecDNAs to accumulate, as shown by our *ex vivo* experiments. In this regard, it will be important to determine whether tissue damage and regeneration can facilitate this process and if tissues with high proliferation rates (intestine, hematopoietic system) will be more prone to ecDNA-driven tumorigenesis. We expect that the tools and the genetically engineered mouse models described here will enable the scientific community to address these and other key questions concerning the biology of ecDNAs and their roles in tumor initiation and tumor progression.

## Methods

### Plasmids and viral vectors

Plasmids containing the “circularization cassettes” were generated by using NEBuilder HiFi DNA Assembly Cloning Kit (#E5520S; New England BioLabs) and KLD Enzyme Mix (#M0554S; New England BioLabs) and validated by Sanger sequencing. Purified, high titer recombinant adenoviruses encoding Cre (AdCre) were purchased from Viraquest (VQ-Ad-CMV-Cre; 1×10^12^ particles/ml, #091317; ViraQuest) and University of Iowa (Ad5CAGCre; VVC-U of Iowa-8193, University of Iowa). pBABE-Puro and pBABE-Puro-HRAS^G12V^ plasmid were obtained from Addgene. For retrovirus production, HEK-293T cells were seeded in DMEM high-glucose supplemented with 10% FBS without antibiotics in 100 mm petri dishes. The next day, cells at approximately 60-70% confluence were transfected with 20 µg of retrovirus carrying *HRAS^G12V^* cDNA or a control vector, 5 µg of packaging plasmid, and 1 µg of envelope plasmid. After 24 hours, the medium was replaced with DMEM medium with antibiotics. Cells were incubated for 48 hours with two consecutive harvesting of the medium containing the retroviral particles at 24 and 48 hours. Collected medium upon 24 and 48 hours was filtered using a 0.45 µm filter unit and used to transduce MEF cells. Retroviral infection was performed incubating cells with viral supernatant supplemented with polybrene (0.2 µl/ml, #TR-1003G, Millipore Sigma).

### Cell Culture

Culture medium of HCT116 cells (ATCC, CCL-247) was McCoy’s 5a Medium Modified (#16600108; Thermo Fisher Scientific) supplemented with 10% heat inactivated FBS (#F2442; Sigma Aldrich), penicillin/streptomycin (100 U/l, #400-109; Gemini Bio-Products). For selection of targeted cells engineered with the “circularization cassettes”, the following antibiotics were used: puromycin (2 μg/ml, #P9620; Sigma Aldrich), hygromycin B (200 μg/ml, #10687010; Thermo Fisher Scientific). HEK-293T cells for viral vectors production were cultured in DMEM high-glucose (w/ 4.5 g/l; #11995065; Thermo Fisher Scientific) with 10% heat inactivated FBS, L-Glutamine (2 mM), and penicillin/streptomycin (100 U/l).

Mouse embryonic fibroblast (MEF) with different genotypes were isolated from E13.5 mouse embryos and propagated in DMEM high-glucose supplemented with 10% heat inactivated FBS, L-Glutamine (2 mM), and penicillin/streptomycin (100 U/l).

Adult neural Stem Cells (aNSC) were isolated and cultured following the protocol described in Ahmed et al. ^39^. Upon isolation, aNSC were maintained as neurosphere and then allowed to attach to laminin-coated (#L2020; Sigma Aldrich) dishes in Neurocult Stem Cell Basal Media with NeuroCult Proliferation Supplement (Mouse & Rat) (#05702; Stem Cells Technologies), 20 ng/ml EGF (#78006; Stem Cells Technologies), 10 ng/ml bFGF (#78003; Stem Cells Technologies), and 2 μg/ml heparin (#07980, Stem Cell Technologies). All cells were tested for mycoplasma contamination. Cells were maintained in a humidified, 5% CO2 atmosphere at 37 °C

### Focus formation assay

For the focus formation assay, MEFs transduced with pBABE-Puro or pBabe-Puro-HRASG12V were selected in puromycin for 72 hours and plated at ∼100,000/well into 6 well plates. Medium was changed every 4 days. After 2 weeks wells were fixed in 4% paraformaldehyde, stained with Giemsa, washed in distilled water and photographed.

### In vivo tumor formation

For *in vivo* tumor formation assay, nude mice were injected in the flank with 400,000-1,000,000 pBABE-Puro-HRASG12V transduced MEFs of the indicated genotype. Mice were monitored every 2-3 days and euthanized when the tumor volume reached 2 cc.

### Homologous recombination in HCT116 cells

Cas9 protein, crRNA, and tracrRNA were purchased from Integrated DNA Technologies pre-assembled by incubating according to manufacturer’s instructions.

The two circularization cassettes were introduced sequentially. The donor cassettes were amplified using primers containing 80 nucleotides 5’ homology sequence to the desired targeting site. 100-500 ng of gel-purified PCR products were transfected into 500,000 HCT116 cells plated the day before in a well of a 6-wells plate using Lipofectamine 3000 (Thermo Fisher). Two hours later, the pre-assembled Cas9-crRNA-tracrRNA were introduced using the Lipofectamine CRISPR-Max (Thermo Fisher) following manufacturers instructions. Selection with either puromycin (5’ cassette, 2µg/ml) or hygromycin (3’ cassette, 200µg/ml) was started 48 hours after transfection. Surviving clones were isolated and screened by PCR followed by Sanger sequencing to detect correct targeting.

### Animals

All animal experiments were approved by MSKCC’s Institutional Animal Care and Use Committee. To generate the *Myc^ec^* and *Mdm2^ec^* mouse strains in a C57BL/6J background, zygote electroporation was carried out by the MSKCC Mouse Genetics Core Facility based on published protocols^40^ by using crRNAs experimentally validated in mouse ESCs. Briefly, multiple zygotes placed in an electrode chamber were subjected to electroporation at one time. Each electroporation mixture contained the 5’ and 3’ breakpoint targeting crRNAs (25 ng/μl), *S.p.* Cas9 V3 protein (100 ng/μl; IDT), and two ssODNs of 159 bp with asymmetric homology arms (200 ng/μl) in a solution of 0.1% polyvinyl alcohol. Electroporated zygotes were cultured in KSOM medium at 37°C and 5% CO_2_ until the two-cell stage, after that they were transferred to the oviducts of pseudo-pregnant females on the day of the vaginal plug. N0 animals generated from the zygotes were genotyped and Sanger sequenced to confirm insertion of both *loxP* sites. Double-targeted N0 mice were mated to C57BL/6J wild type mice to generate *Myc^ec/+^* F1 progeny. Primers for genotyping are listed in **Supplementary Table 2**. The *Mdm2^ec^*mouse was also generated as described above except that gRNAs were used to target chr10:116711442 (TCTTACAGCATACTACGGTC TGG) and chr10:118002454 (TTCTGCGATTCGTTATGCGT AGG) for the 5’ and 3’ *loxP* insertion sites, respectively.

### Pathology

Tumors were fixed in formalin overnight, embedded, and sections were stained with hematoxylin and eosin. Slides were reviewed by a C.R.A. a board-certified pathologist with extensive expertise in the characterization of human sarcomas and liposarcomas.

### Antibodies, immunoblots

Cells were lysed in Laemmli buffer, supplemented with protease (cOmplete™, #COEDTAF-RO; Roche) and phosphatase inhibitors (EDTA-free Protease Inhibitor cocktail; #PHOSS-RO; Roche). Proteins were separated in SDS-PAGE and analyzed by western blotting by standard procedures. After protein transfer, the nitrocellulose membranes (#1704271; BioRad) were blocked by incubation with LICOR Intercept (TBS) blocking buffer. The following primary antibodies were used: anti-VINCULIN (1:5,000; Millipore, #MAB3574), anti-c-MYC (1:1,000; Cell SIgnaling, #D84C12), anti-MDM2 (1:1000, Cell Signaling, #51541), anti-alpha-tubulin (1:5000, Cell Signaling, #3873), anti-p53 (1:1000, Leica Biosystems, P53-PROTEIN-CM5), anti-p21 (1:1000, Cell Signaling, #64016). The following secondary antibodies were used: IRDye 800 anti-Rabbit (#926-32213, LICOR) and IRDye 680 anti-Mouse (#926-68072, LICOR). Immunostained bands were detected by using an Odyssey Imaging System (LICOR). H&E slides were examined by C.A. a board-certified pathologist with extensive expertise in human sarcomas and liposarcomas.

### RNA extraction and RT-qPCR

Total RNA was isolated using the RNeasy Mini Kit (QIAGEN, #74106) according to the manufacturer’s instructions. After treatment with DNAse I (Ambion, #AM2222), 1 μg of purified RNA was retro-transcribed with oligos d(T)_18_ by using SuperScript IV First-strand System (Invitrogen, #18091050). For RT-qPCR, an aliquot of the RT reaction was analyzed with PowerUp SYBR Green (Applied Biosystems, #A25777) a QuantStudio™ 6 Flex real-time PCR system (Applied Biosystems). Target transcript levels were normalized to those of the indicated reference genes. The expression of each gene was measured in at least three independent experiments.

### Milademetan dose-response curve

A total of 1,500 cells were seeded in each well of a 96-well plate, with 100 µl of complete media per well. The cells were allowed to adhere for 12 hours in the incubator before initiating the treatment. Milademetan (MedChemExpress, HY-101266) doses ranging from 20 to 10,240 nM were prepared, along with a vehicle control at the highest treatment dose. The doses were added to the respective wells in a volume of 100 µl, ensuring that the cells received Milademetan doses ranging from 10 to 5120 nM. Following the addition of the treatment doses, the cells were incubated under standard growth conditions for a total of 72 hours to allow the treatment to exert its effects.

After the 72-hour incubation period, cell viability was determined using the Cell-TiterGlo assay (Promega, cat # G7570, following manufacturer’s instructions. Briefly, the assay reagent was added to the wells, and the plates were gently agitated to ensure thorough mixing. The luminescence signal was then measured using a Synergy 2 plate reader (Biotek) according to the manufacturer’s guidelines.e

### Gene Copy Number Assay

*Myc*, *MDM2,* and *Mdm2* gene amplification was evaluated by using TaqMan™ Copy Number Assays (Probe Mm00734221_cn, Probe Hs02873318_cn and Probe Mm00312030_cn) by using a QuantStudio™ 6 Flex real-time PCR system. Briefly, genomic DNA was amplified by using the TaqPath™ ProAmp™ Master Mix (Applied Biosystems, #A30865) kit and following supplier’s instructions. TaqMan™ Copy Number Reference Assay (Tfrc, #4458367 and TERT, #4403316) were used in duplex as reference for mouse and human genomes, respectively. Relative copy number variations were calculated by using the ddCt method.

### RNAseq and GSEA analysis

Paired ends RNAs libraries were sequenced by the MSKCC integrated genomics core. Reads were mapped to mm10 using the *STAR* aligner ^41^ and differential gene expression was calculated using the *DESeq2* R package ^42^. For GSEA, genes were ranked based on their moderated p-value [-log10(adjPavl)*sign(log2foldchange)], gene sets were obtained from the HALLMARK msigdb pathways database. Enrichment was calculated using the *fgsea* R package.

### Shallow Whole-Genome Sequencing (sWGS)

Shallow Whole-Genome Sequencing was carried out by the Integrated Genomics Operation (IGO) core at MSKCC. Briefly, after PicoGreen quantification and quality control by Agilent BioAnalyzer, 100ng of genomic DNA was sheared using a LE220-plus Focused-ultrasonicator (#500560; Covaris) and sequencing libraries were prepared using the KAPA Hyper Prep Kit (#KK8504; Kapa Biosystems) with 8 cycles of PCR. Samples were run on a NovaSeq 6000 in a PE100 run, using NovaSeq 6000 S4 Reagent Kit v1.5 (200 cycles) (Illumina). Paired ends genomic DNA libraries were sequenced by the MSKCC integrated genomics core. Reads were mapped to mm10 or hg19 using the Bowtie2 aligner ^43^. To calculate genome coverage and copy number changes we used the QDNAseq R package ^44^ with 15kb bins. Plots were generated using the Gviz package ^45^.

### Amplicon Architect Analysis

We utilized the AmpliconSuite-pipeline (version 0.1555.1, https://github.com/AmpliconSuite/AmpliconSuite-pipeline) to invoke AmpliconArchitect ^46^(version 1.3.r5) on a collection of copy-number seed regions generated using CNVkit ^47^(version 0.9.9), with default settings.

### Metaphase Chromosome Spread analysis

Cells were incubated with KaryoMAX (#15212012; Thermo Fisher Scientific) treatment at 0.05 μg ml^!"^ for 1 h (mouse cells) or at 0.1 μg ml^!"^ for 1h and 30 min (human cells). Single-cell suspension was then collected, washed in PBS, and treated with 75 mM KCl for 10 min at 37 C. Samples were then fixed in ice-cold 3:1 methanol:glacial acetic acid (Carnoy’s fixative) for 20 min and washed an additional three times with the Carnoy’s fixative. Fixed cells were dropped onto glass slide in a humidified chambers and counterstained with DAPI (#H-1800 Vector Laboratories). Images were acquired with a AX10 Imager.Z1 Zeiss microscope through 63X objective lens. Zeiss Zen 2.3 pro software was used for image acquisition. Fiji (version 2.0.0-rc-65/1.15w) was used for image analysis and for brightness and contrast adjustments. Fiji’s “Invert LUT” and “Shadows” postprocessing were sequentially applied to better visualize double minutes.

### Fluorescence *In Situ* Hybridization (FISH) analysis

DNA FISH analysis was performed on fixed cells using a 2-color probe. *MDM2* (Green dUTP) and centromeric control (orange dUTP) FISH probes were purchased from Empire Genomics (Williamsville, NY) (SKU MDM2-CHR12). BAC clones containing murine *Myc* locus (RP23-295E4, RP23-263C13) were labeled with Red dUTP and RP23-173M6 (11qA1), labeled with Green dUTP served as the control. All RP11 and RP23 clones were purchased from the Roswell Park Cancer Institute Genomics Shared Resource (Buffalo, NY).

Probe labeling, hybridization, post-hybridization washing, and fluorescence detection were performed according to procedures established at the Molecular Cytogenetics Core Facility. Slides were scanned using a Zeiss Axioplan 2i epifluorescence microscope (Carl Zeiss Microscopy, Thornwood, NY) equipped with Isis imaging software (MetaSystems Group Inc, Waltham, MA) or Leica SP5 confocal microscope (Leica) with a 63x objective. The entire hybridized area was scanned through 63X objective lens to assess the quality of hybridization and signal pattern. For each cell line, a minimum of 50 intact nuclei and 15 metaphase spreads were analyzed and signal pattern recorded. To the extent possible, apoptotic cells/bodies were excluded.

The BAC clone RP23-428D5 (BACPAC, Emeryville, CA), was used for the detection of murine *Mdm2* locus. Following inoculation, bacteria cells were pelleted and BAC DNA was extracted using the NucleoBond Xtra BAC kit (Takara, 740436) as per manufacturer’s instructions. The DNA was then amplified using the BioPrime DNA Labeling System (Thermo, 18094011) and labeled with ChromaTide Alexa Fluor 568-5-dUTP (Thermo, C11399) as per manufacturer’s instructions. The labeled probe was precipitated with isopropanol and stored in 70% ethanol in - 20C indefinitely. Prior to hybridization, the labeled probe was pre-annealed with mouse COT-1 DNA in hybridization buffer (2xSSC, 50% formamide, 10% dextran sulfate, 0.1% SDS) for 90 minutes at 37 °C. Hybridization of slides using pre-annealed probes was performed at 72C for 2 minutes followed by 37 °C overnight. Post-hybridization washes were conducted in 0.4xSSC/0.3% Igepal at 72 °C for 2 minutes followed by 2xSSC at room temperature for 5 minutes. Slides were then rinsed briefly in water, air dried, counter-stained with DAPI (Thermo, D1306), and mounted with Prolong Diamond Antifade Mountant (Thermo, P36965). Images were acquired in the MSKCC Molecular Cytology Core using a Zeiss Imager equipped with a Zeiss AxioCam Mrm camera and a 100x oil objective.

*Mdm2* RNA FISH was performed in collaboration with the MSKCC Molecular Cytology Core. Briefly, paraffin-embedded tissue sections were cut at 5 μm and kept at 4 °C. Samples were loaded into Leica Bond RX autostainer, baked for 30 minutes at 60 °C, dewaxed with Bond Dewax Solution (Leica, AR9222), and pretreated with EDTA-based epitope retrieval ER2 solution (Leica, AR9640) for 15 minutes at 95°C. The mouse MDM2 probe (Advanced Cell Diagnostics (ACD), Cat# 447648) was hybridized for 2 hours at 42 °C. Mouse PPIB (ACD, Cat# 313918) and dapB (ACD, Cat# 312038) probes were used as positive and negative controls, respectively. The hybridized probes were detected using RNAscope 2.5 LS Reagent Kit – Brown (ACD, Cat# 322100) according to manufacturer’s instructions with the following modifications: DAB application was omitted and replaced with Fluorescent CF594/Tyramide (Biotium, B40953) for 20 minutes at RT. Images were acquired using a Zeiss Imager equipped with a Zeiss AxioCam Mrm camera, a 20x air objective, and a 100x oil objective.

### Statistic and reproducibility

Paired or unpaired Student’s two-tailed t-test and one-way ANOVA (corrected for multiple comparisons, Turkey test), were used to compare two or more groups, respectively, and to determine statistical significance (GraphPad Prism 9) Welch’s correction was used for population with unequal variances. When not differently indicated the mean value and the standard deviation of each condition are shown. Differences were considered significant at P value <0.05.

## Acknowledgements

We thank the group of Gigi Lozano for discussion and advice on the biology of Mdm2 and for sharing the Mdm2^−/−^; p53^−/−^ double KO MEFs that were used to validate antibodies for Western Blot. This work was supported by grants from NIH-NCI (P30 CA008748), the American Cancer Society (Discovery Boost grant), and by a rapid response grant from the Functional Genomics Initiative. DP was supported by an AIRC fellowship for Abroad, MZ is a recipient of the Paul Calabresi Career Development Award for Clinical Oncology from the Memorial Sloan Kettering K12 Clinical Translational Cancer Research Training Program, MW is supported by the Erwin Schrodinger Fellowship from the Austrian Science Fund (FWF, J4723), MY is supported by the Harold E. Varmus Fellowship in Cancer Biology. KMG was supported by ETIUDA scholarship (No. 2018/28/T/NZ5/00467) funded by the National Science Center of Poland. We thank members of the Benezra, the Joyner, and the Lowe laboratories for helpful discussion and advice.

## Author contributions

A.V. conceived the project and designed the general strategy to generate ecDNAs *in vitro* and *in vivo*. R.G., M.Y., F.G., and A.V. generated and tested the circularization cassettes; M.Y., K.M.G., T.M., D.P., M.L. and A.V. generated and characterized the HCT116 *invMDM2* and *ecMDM2* cell lines. R.G. generated the *Myc^ec^* mouse; D.P. characterized the *Myc^ec^* mouse and performed the experiments in adult neural stem cells, with assistance from H.M.H.N. and M.L.; M.Z. generated the *Mdm2^ec^* mouse and, together with A.V., M.Y. and M.W. characterized it and studied the effects of ecDNA formation in MEFs. D.P., generated the RNAseq data and, together with A.V., analyzed them. A.V. analyzed the sWGS datasets; Y.C.F. tested the effects of Milademetan on *Mdm2^ec/+^* transformed MEFs and performed the RT-PCR, G.L.R., and Z.J performed the western blots. C.R.A. analyzed the tumors generated from *Mdm2^ec/+^* cells; J.L. and V.B. performed the Amplicon Architect analysis. C.M. and K.C. genotyped and maintained the mouse strains. D.P., M.Z., and A.V. wrote the manuscript with assistance and comments from all co-authors.

## Materials & Correspondence

Requests for materials and reagents should be directed to AV

## Supplementary Figures Legend

**Supplementary Figure 1.**
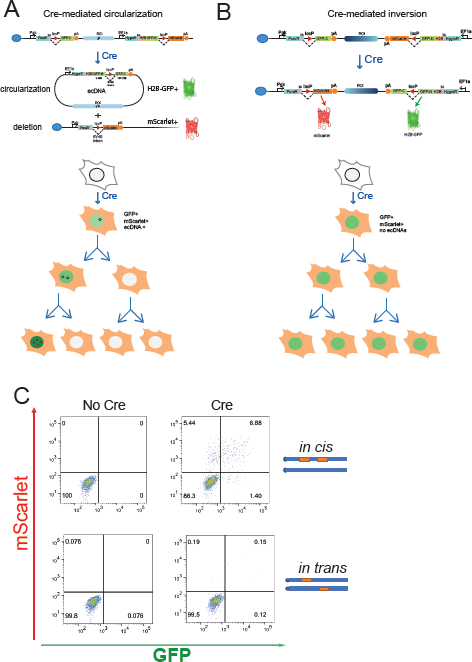
**A.** Schematic and predicted outcomes of the circularization strategy. Upon Cre-mediated recombination, ecDNAs encoding GFP are generated, while mScarlet is expressed from the linear chromosome harboring the corresponding deletion. Random segregation of the ecDNAs will lead some cells to acquire extra copies of the ecDNAs and therefore become more strongly positive for GFP, while other cells will lose the ecDNAs and become GFP-negative. **B.** Schematic and predicted outcome of the inversion strategy. In this allele, the 3’ circularization cassette is inserted at the same location as in the *ecDNA* allele, but with opposite orientation. Upon Cre-mediated recombination, the entire region is inverted resulting in the expression of both GFP and mScarlet reporters. Because both reporters remain on the chromosome, the resulting cells are predicted to remain double positive for mScarlet and GFP indefinitely. **C.** Discrimination between *in cis* and *in trans* cassette integration. Representative flow cytometry analysis of two control or AdCre-infected *ecMDM2* HCT116 clones targeted with the two circularization cassettes examined 6 days after infection. For both clones, the correct integration and sequence of each cassette was confirmed by Sanger sequencing, however the clone shown on top displays significantly higher recombination efficiency, indicating *in cis* integration of the *loxP* sites.

**Supplementary Figure 2.**
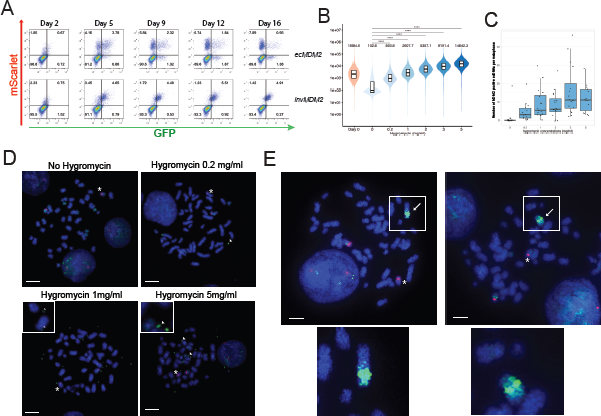
**A.** *ecMDM2* and *invMDM2* cells were infected with AdCre, expanded in the absence of hygromycin, and analyzed by flow cytometry at the indicated timepoints. Clones were propagated for 6 days in the presence or absence of hygromycin (200µg/ml) and analyzed by flow cytometry. Pseudocolor scatter plots of GFP and mScarlet fluorescence at each time point are shown. Notice the progressive disappearance of double positive cells and the concomitant increase in GFP-;mScarlet+ cells in the *ecMDM2* samples. **B.** Violin plots showing GFP intensity of sorted double positive *ecMDM2* and *invMDM2* cells expanded in the presence of the indicated concentration of hygromycin for 13 days and analyzed by flow cytometry (see also figure 2F-H). Median GFP intensity for each sample is also indicated. Boxes indicated interquartile range. **** indicates p-value < 0.0001 as determined by the Wilcoxon–Mann–Whitney test**. C.** *ecMDM2* and *invMDM2* cells were infected with AdCre and double positive cells were sorted and expanded in the presence of the indicated amount of hygromycin for 13 days and analyzed by DNA-FISH using an MDM2 probe (see also Fig. 2F). Shown are box-and-whisker plots of *MDM2*-containing ecDNAs per metaphase for cells incubated in the indicated amount of hygromycin. **D.** Representative metaphase spreads containing *MDM2-*positive ecDNA for the indicated hygromycin concentration as quantified in C. Scale bar: 7.5 µm. **E.** Presence of *MDM2-*positive HSRs in representative metaphase spreads. Scale bar: 7.5 µm.

**Supplementary Figure 3.**
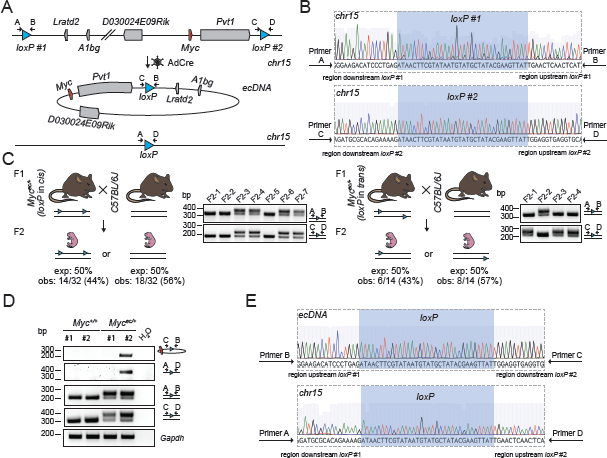
**A.** Schematic of the *Myc^ec^* allele. Upon AdCre transduction, Cre recombinase promotes the excision and circularization of the genomic region flanked by the two *loxP* sites. Genes are indicated in gray boxes. *Myc* gene is highlighted in red. Arrows indicate primers to detect the insertion of *loxP* sites for genotyping and Sanger sequencing. **B.** Sanger sequencing results of inserted *loxP* sites in the F1 progeny. *LoxP* sequences in the correct orientation are highlighted in yellow. **C**. Breeding schemes to test the insertion of *loxP* sites in the same chromosome – *in cis* – (left) or in different homologous chromosomes – *in trans* – (right) are indicated. Expected and observed genotypes in the F2 generation are reported. Representative genotyping PCR results are also shown. **D.** gDNA-PCR analysis showing the formation of a neo-junction (Primers C-B) on the circularized genomic region, the deletion (Primers A-D) on the linear chromosome, the insertion of *loxP* sites (Primers A-B and C-D) in wild type (*Myc^+/+^*) and *Myc^ec/+^*aNSC transduced with AdCre. *Gapdh* as a control. Three different MEF are shown. **E.** Sanger sequencing results of reconstituted *loxP* sites in the ecDNA (top) and in the *Myc* locus on chromosome 15 (bottom).

**Supplementary Figure 4.**
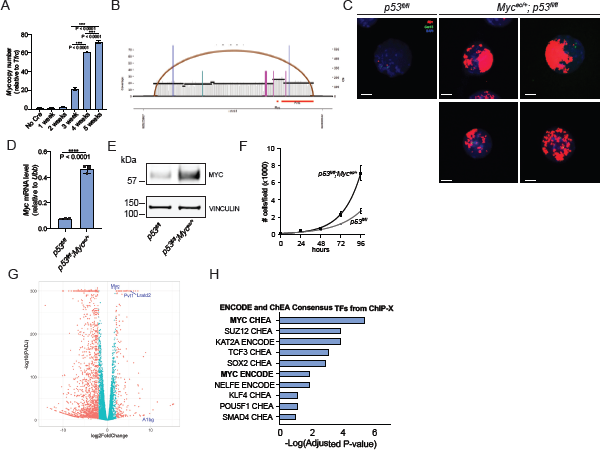
**A.** Myc gene relative copy number ecDNAs in *Myc^ec/+^; p53^fl/fl^* aNSC at different time points upon AdCre infection. Error bars indicate SD. one-way ANOVA for multiple comparisons. P-value is indicated. **** < 0.0001. **B.** sWGS data from the 5 week time point (see Figure 3B) were analyzed using AmpliconArchitect to identify structural variants. The coverage and structural variant plot reveals a structural variant closing the left-and-right endpoints of the amplified region forming an ecDNA-like cycle spanning the region flanked by the *loxP* sites. **C.** Representative interphase-nuclei FISH (right) on aNSC from *Myc^ec/+^; p53^fl/fl^* 5 weeks upon AdCre infection showing clustering inside the nucleus. *Myc* probe is in red. Control probe labeling a pericentromeric region of chromosome 15 is in green. Scale bar: 5 µm. **D.** Analysis, by RT-qPCR, of *Myc* expression levels (relative to *Ubb*) in *Myc^ec/+^; p53^fl/fl^* and *p53^fl/fl^* aNSC treated with AdCre and propagated for 5 weeks to allow ecDNA accumulation. n = 3 biological replicates. Unpaired two-tailed t test. P-value < 0.0001. Error bars indicate ±SD. **E**. Immunoblotting of MYC expression levels in *Myc^ec/+^; p53^fl/fl^* and *p53^fl/fl^* aNSC 5 weeks upon AdCre Infection. VINCULIN as loading control. **F**. Growth curves of *Myc^ec/+^;p53^fl/fl^*;(black) and control *p53^fl/fl^*(grey) aNSC 5 weeks after AdCre infection. n = 10 biological replicates, 3 fields for each replicate have been acquired. **G**. Volcano plot of DEGs in *Myc^ec/+^; p53^fl/fl^*aNSC 5 weeks upon AdCre infection. Genes with log2 Fold Change > 1, and adjusted P-value < 0.01 are indicated in red. Genes located in the *Myc^ec^* amplicon are labeled. **H**. Enrichment analysis of DEGs in *Myc^ec/+^; p53^fl/fl^* aNSC for putative transcription factor binding sites performed by using EnrichR webtool and using the ChEA and ENCODE gene-set datasets.

**Supplementary Figure 5.**
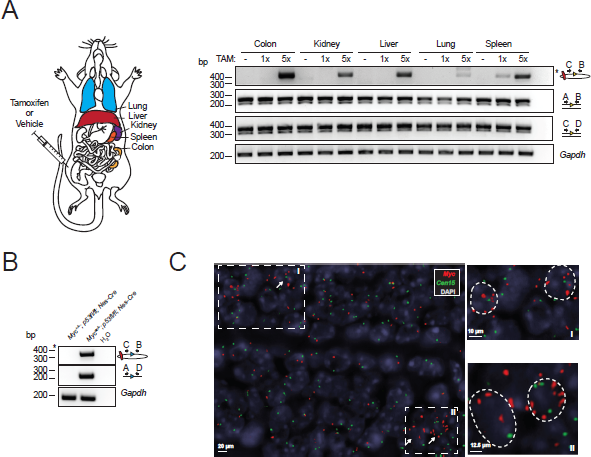
**A**. gDNA-PCR analysis with primers design to detect the circularized allele upon Cre-expression (Primers C-B) performed on DNA extracted from the indicated organs collected from *Myc^ec/+^; p53^fl/fl^*; Rosa26-Cre-ER^T2^treated 0, 1, or 5 times with tamoxifen. *Gapdh* is included as a PCR control. **B**. gDNA-PCR analysis showing circularization (Primers C-B) and deletion (Primers A-D) of the region flanked by the *loxP* sites in brain collected from *Myc^+/+^* and *Myc^ec/+^*; *p53^fl/f^*; Nestin-Cre adult mice (4 months old). **C**. DNA FISH on brain tissue slide from *Myc^ec/+^; p53^fl/f^*; Nestin-Cre adult mice (4 months old). The subventricular zone (SVZ) is shown. *Myc* probe is in red. Control probe labeling a pericentromeric region of chromosome 15 is in green. Insets are high magnification showing multiple *Myc* signals per nucleus are on the right.

**Supplementary Figure 6.**
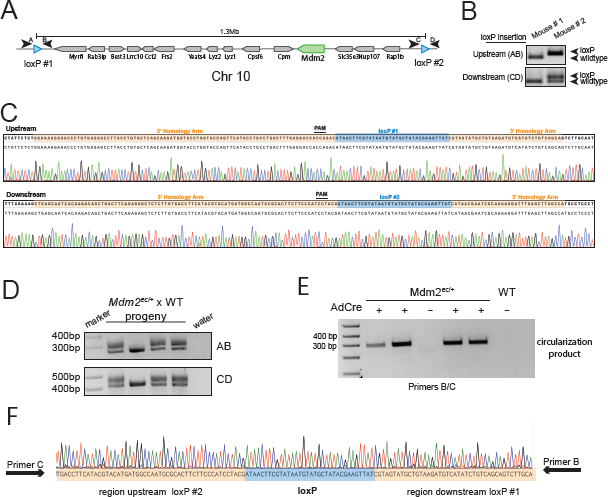
**A.** Schematic of the insertion of two *loxP* sites flanking the mouse *Mdm2* locus, spanning a total of 1.3Mb. Arrowheads indicate the PCR primers used to identify successful *loxP* insertion. **B.** Zygotes were injected with Cas9-gRNA complexes and donor DNA containing *loxP* sites. Genotyping of F0 mice shows that Mouse #2 has *loxP* sites inserted at both the upstream and downstream locations. Note that the upstream wildtype band for Mouse #2 is lost, in this case indicating homozygous insertion of the *loxP*. **C.** Chromatograms showing correct insertion of the two *loxP* sites. **D.** A heterozygote F1 progeny of Mouse #2 crossed to a wildtype mouse shows that the loxP bands for the upstream and downstream integration sites co-segregate in the resulting F2 progeny, indicating that the *loxP* sites were inserted *in cis*. **E.** AdCre-treated *Mdm2^ec/+^* MEFs develop recombination of the *loxP* sites, resulting in the circularization of the intervening region as indicated by the amplification product of primers B and C. **F.** Chromatogram obtained by Sanger sequencing of the circularization product from E) demonstrated expected recombination sequence surrounding the *loxP* site.

**Supplementary Figure 7.**
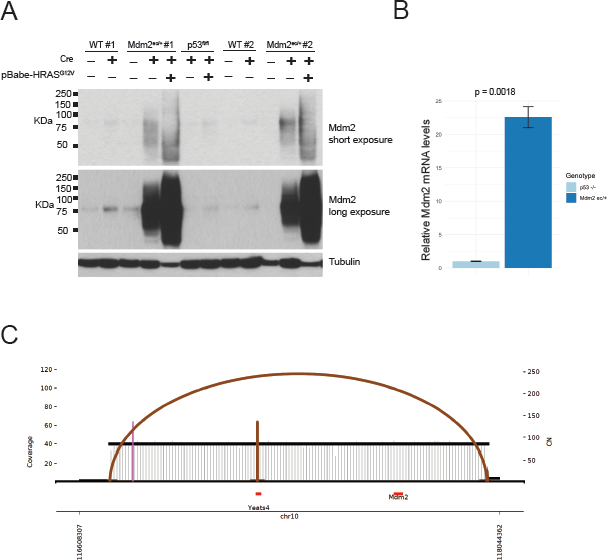
**A.** Western blot for MDM2 and tubulin of wildtype, *Mdm2^ec/+^*, and *p53^fl/fl^* MEFs with or without treatment by Cre and *HRAS^G12V^*show significant accumulation of Mdm2 only in Cre-treated *Mdm2^ec/+^*MEFs. **B.** qPCR in Cre-treated *Mdm2^ec/+^*, and *p53^fl/fl^*MEFs show preferential upregulation of *Mdm2* transcripts in *Mdm2^ec/+^*MEFs. **C.** sWGS data from AdCre- and HRAS infected *Mdm2^ec/+^* MEFs (see Figure 4) were analyzed using AmpliconArchitect to identify structural variants. The structural variant plot reveals a structural variant closing the left-and-right endpoints of the amplified region forming an ecDNA-like cycle spanning the region flanked by the *loxP* sites.

**Supplementary Figure 8.**
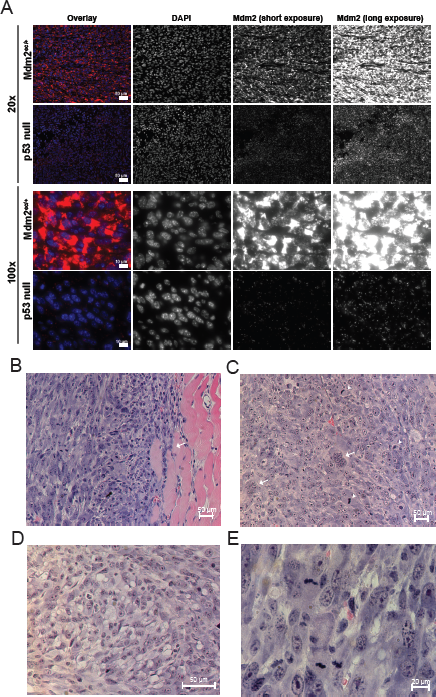
**A.** RNA-FISH using an *Mdm2* probe on tumor tissues from *Mdm2^ec/+^;HRAS^G12V^* and *p53^fl/fl^;HRAS^G12V^*MEFs at both low (20x, scale bar = 50µm) and high (100x, scale bar = 10µm) magnifications. **B.** Low magnification view showing a high-grade spindle cell sarcoma arranged in short fascicles and infiltrating into skeletal muscle. **C.** The lesional cells show increased nuclear pleomorphism, with scattered multinucleated forms (arrows) and increased mitotic activity (arrowheads). **D.** Higher magnification shows solid sheets of epithelioid to ovoid cells with distinct single or multi-intra-cytoplasmic fat vacuoles consistent with signet ring lipoblasts. Focal nuclear indentation, a characteristic feature of lipoblasts is also noted. **E.** Increased mitotic activity and pleomorphic spindle cells with amphophilic cytoplasm and ovoid nuclei with clumped chromatin and prominent nucleoli in keeping with a high grade sarcoma.

**Supplementary Figure 9.**
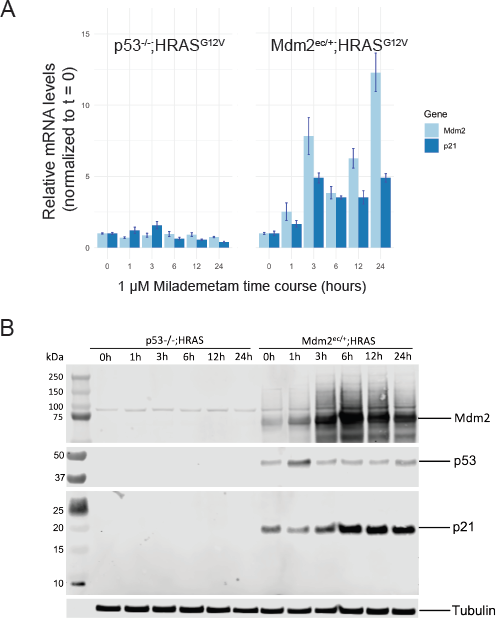
Cre-treated *Mdm2^ec/+^;HRAS^G12V^*and *p53^fl/fl^;HRAS^G12V^*MEFs were exposed to 1µM milademetan, collected at indicated time points, and analyzed by RT-qPCR (**A**), and immunoblot (**B**). Mdm2 and p21 mRNA and protein products are rapidly and strongly induced selectively in the *ecMDM2* cells in response to milademetan.

## Supplementary Tables

***Supplementary Table 1*** RNASeq differential gene expression in *Myc^ec/+^;p53^fl/fl^* vs *p53^fl/fl^* aNSCs cells 5 weeks after AdCre infection.

***Supplementary Table 2*** Oligonucleotides, gRNAs, and PCR primers used in this study.

